# Antibiotic Resistance Genes and Taxa Analysis from Mat and Planktonic Microbiomes of Antarctic Perennial Ice-covered Lake Fryxell and Lake Bonney

**DOI:** 10.1101/2021.12.17.473166

**Authors:** Sheetal Tallada, Grant Hall, Daniel Barich, Rachael M. Morgan-Kiss, Joan L. Slonczewski

## Abstract

The perennial ice-covered lakes of the Antarctic McMurdo Dry Valleys harbor oligotrophic microbial communities that are separated geographically from other aquatic systems. Their microbiomes include planktonic microbes as well as lift-off mat communities that emerge from the ice. We used ShortBRED to quantify the antibiotic resistance genes (ARGs) from metagenomes of lift-off mats emerging from ice, and from filtered water samples of Lake Fryxell and Lake Bonney. The overall proportion of ARG hits was similar to that found in temperate-zone rural water bodies with moderate human inputs. Specific ARGs showed distinct distributions for the two lakes, and for mat versus planktonic sources. Metagenomic taxa distributions showed that mat phototrophs consisted mainly of cyanobacteria or betaproteobacteria, whereas the water column phototrophs were mainly protists. An enrichment culture of the betaproteobacterium *Rhodoferax antarcticus* from a Lake Fryxell mat sample showed an unusual mat-forming phenotype not previously reported for this species. Its genome showed no ARGs associated with betaproteobacteria, but had ARGs consistent with a minor *Pseudomonas* component. The Antarctic lake mats and water showed specific ARGs distinctive to the mat and water sources, but overall ARG levels were similar to those of temperate water bodies with moderate human inputs.

## INTRODUCTION

One of Earth’s coldest dry deserts, the McMurdo Dry Valleys in Antarctica possess a string of stratified lakes with unique geochemistry and microbial communities (Spigel & Priscu 1998, Roberts *et al*. 2004, Cavicchioli 2015, Sohm *et al*. 2020). The microbial genomic analysis of these lakes remains limited (Dillon *et al*. 2020, Koo *et al*. 2018, Wei Li *et al*. 2020). There are a number of reports on antibiotic resistance genes (ARGs) in Antarctic soil or marine bacteria (Antelo *et al*. 2021, Tam *et al*. 2015, Wang *et al*. 2016, Na *et al*. 2019, van Goethem *et al*. 2018) but few studies of ARGs in Antarctic lake water or their mat communities (Jara *et al*. 2020). The prevalence of ARGs in relatively pristine habitats is of compelling importance for human health and environment, as it provides a baseline for assessment of human-associated ARG contamination (Allen *et al*. 2010, D’Costa *et al*. 2011). Antimicrobial resistance is a leading cause of death worldwide (Murray *et al*. 2022).

We report on the taxonomic composition and ARG prevalence in the water of Lake Fryxell and Lake Bonney, and in the phototrophic mat communities of Lake Fryxell (**Fig. 1A**). Lake Fryxell extends along the Taylor Valley, 10 km downstream of Lake Bonney, 80 km across the Ross Sea from McMurdo Station. No permanent stream connects the two lakes; their main hydrology involves glacial meltwater, and each lake has an underground brine aquifer (Mikucki *et al*. 2015). Their water ecosystems are entirely microscopic, with no fauna larger than nematodes. The lakes and their ecosystems are studied as models for ancient Mars (Head & Marchant 2014) and as indicators of climate change (Hall *et al*. 2017).

**Figure 1.**
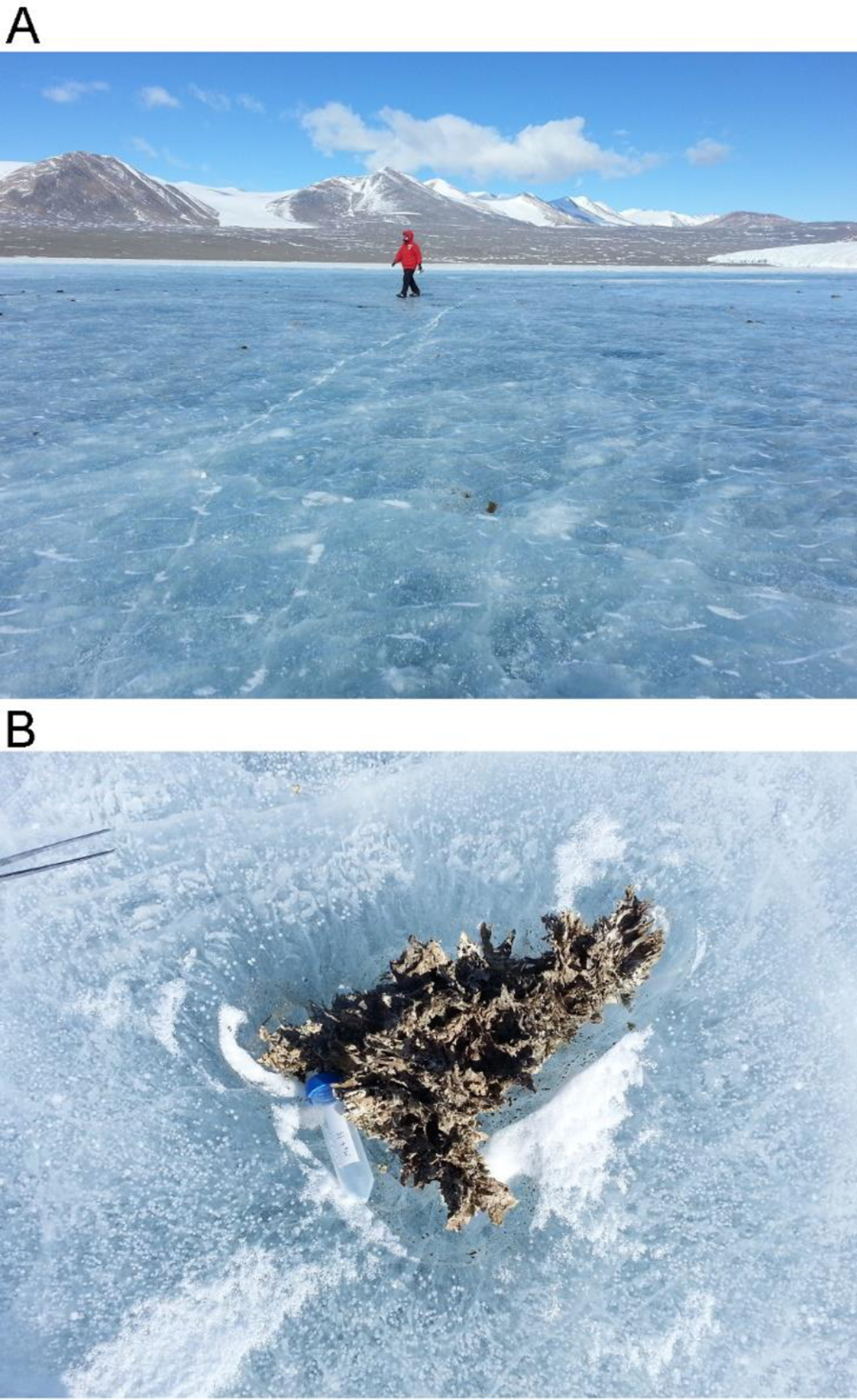
Lake Fryxell uplift mats. (A) Lake Fryxell with permanent ice cover, weathered by katabatic winds; December 10, 2014. (B) Source of MAT-4 DNA. Uplift mat emerges from the ice; approximately 40 cm across.

Lakes Fryxell and Bonney have permanent stratified layers in the water column that give rise to oligotrophic microbial communities (Roberts *et al*. 2004, Li *et al*. 2016, Kwon *et al*. 2017). The lack of turnover allows for the growth of benthic microbial mats that form flame-shaped towers several centimeters tall (Jungblut *et al*. 2016, Hawes *et al*. 2013). The microbial mats have been studied with primary emphasis on the role of cyanobacteria such as *Microcoleus* and *Oscillatoria* (Taton *et al*. 2003, Jungblut *et al*. 2016) although metagenomes reveal a high abundance of proteobacteria (Dillon *et al*. 2020) and protists (Li *et al*. 2016, Bielewicz *et al*. 2011).

Known as “lift-off” mats, the benthic biofilms reach upward, buoyed by oxygen bubbles, until pieces break off and float upward to the undersurface of the ice cover (Parker *et al*. 1982, Moorehead *et al*. 1999). During winter, water freezes beneath the mat fragments, adding to the ice layer, but above the lake the valley’s dry winds ablate the ice. Ice sections reveal 6-8 annual layers of freezing below followed by ablation above (Parker *et al*. 1982) leading to exposure of the dessicated mats (**Fig. 1B**). Surviving several years trapped in ice, the mat material remains viable, and psychrophiles may continue growing (Boetius *et al*. 2015). When mat organisms reach the surface of ablated ice, they can blow off and enter “moats” of melted ice that occur in summer, surrounding the permanent ice in the main part of the lake. The mat material carries communities that include microscopic arthropods and nematodes; thus, the mats can act as vectors for transport of entire ecosystems (Parker *et al*. 1982, Brambilla *et al*. 2001, Dillon *et al*. 2020).

Molecular genetic data on McMurdo Dry Valley microbes remained limited until recently (Taton *et al*. 2003, Vick-Majors *et al*. 2014, Kwon *et al*. 2017). In December, 2014, the austral summer, we obtained water samples from Fryxell and Bonney as well as desiccated mat samples emerging from the ice cover of Fryxell. For the present study, we applied tools of metagenome analysis to compare the taxonomic diversity and ARG abundance of mat and lake samples. We also sequenced the genome of a novel ecotype of *Rhodoferax antarcticus* from a mat-forming enrichment culture, whose phenotype differs markedly from planktonic isolates of this species (Madigan *et al*. 2000, Baker *et al*. 2017, Jung *et al*. 2004).

Our study addressed the important question of ARG prevalence in relatively pristine Antarctic lakes. A modest level of antibiotic resistance is an ancient, widespread phenomenon naturally and historically occurring in all environments (Allen *et al*. 2010, D’Costa *et al*. 2011). Non-anthropogenic processes can select for ARGs in pristine habitats; for example, cyanobacterial blooms drive increases in bacterial ARG prevalence and diversity (Zhang *et al*. 2020). But inputs of human origin can add substantially to the native ARG pool as well as add additional types of ARGs (Antelo *et al*. 2021, Tam *et al*. 2015, Jara *et al*. 2020). We surveyed our mat and water metagenomes for the presence of ARGs, referenced to the Comprehensive Antibiotic Resistance Database (CARD) (Alcock *et al*. 2020).

## METHODS

### Sample collection and culture

Microbial communities were sampled in December, 2014 from two meromictic lakes of the Taylor Valley, Victoria Land, Antarctica. Lake Fryxell has a maximum depth of 20 m (Kwon *et al*. 2017, Lawrence & Hendy 1985), while Lake Bonney has a depth of 40 m (Priscu & Spigel 1996). Both lakes are covered by a perpetual ice layer, approximately 4 m thick (Priscu 2018), although summer melting occurs near the shoreline.

Microbial lift-off mat samples were collected from independent mat tufts emerging separately from the Lake Fryxell ice surface, within the GPS area of: −77.60491, 163.16315; −77.60473, 163.16290; −77.60463, 163.16405; −77.60495, 163.16495. Each sample consisted of a separate tuft of dessicated microbial mat, collected with alcohol-sterilized forceps and stored at −20°C (4 weeks) then at −80°C (indefinitely).

Lake water was sampled from the permanent chemoclines of Lake Fryxell (−77.605, 163.163, 9 m depth) and Lake Bonney, east lobe (−77.719, 162.283, 15 m depth). Samples were obtained using ice holes established by the McMurdo Dry Valleys Long-Term Ecological Research (LTER) program (Priscu 2022b, Priscu 2022a, Priscu 2021b, Priscu 2021a). The LTER geochemical data are presented in Table 1. Water samples were collected in 1-liter cubitainers pre-washed in 10% HCl. Each water sample was concentrated by filtration through a 0.45 µm filter.

**Table 1.**
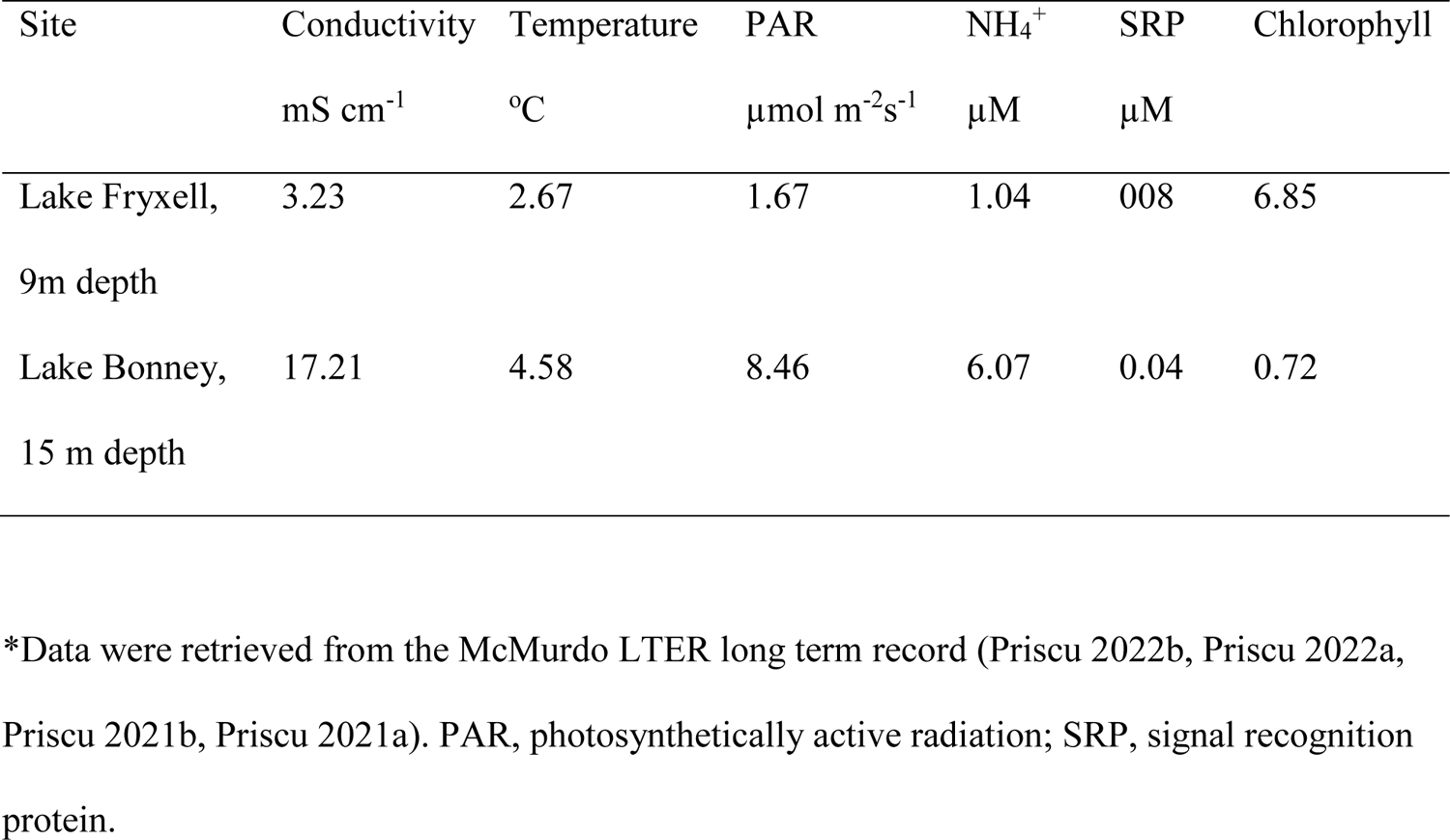
Physical, chemical and biological parameters from Lakes Fryxell and Bonney (east lobe).*

Enrichment culture for anaerobic phototrophs was performed using Harwood PM medium supplemented with 10 mM succinate (Fixen *et al*. 2019, Kim & Harwood 1991, Rey *et al*. 2006). Screwcap pyrex tubes were filled with medium and inoculated with approximately 0.05 g dessicated material from sample Mat-04. Sealed tubes were incubated under approximately 10% PAR at 10°C for five weeks. Portions of biofilm were serially subcultured for two-week periods, then frozen at −80°C. Gram stain was performed by standard methods (Remel kit).

Testing for growth range of pH and NaCl amendment was performed using a *Rhodoferax* medium modified from references (Tayeh & Madigan 1987, Madigan *et al*. 2000). The medium contained: yeast extract (0.5 g/l), EDTA (20 mg/l), malic acid (4 g/l), (NH_4_)_2_SO_4_ (1 g/l), MgSO_4_·7H_2_O (200 mg/l), FeSO_4_·7 H_2_O (12 mg/l), K_2_HPO_4_ (0.9 g/l), KH_2_PO_4_ (0.6 g/l), 10 mM sodium succinate, 100 mM MOPS (3-(N-morpholino)propanesulfonic acid) adjusted to pH 7.0 with KOH. In addition, 1ml/l of a trace elements solution was added [H_3_BO_3_ (2.8 g/l), MnSO_4_·H_2_O (1.6 g/l), Na_2_MoO_4_ ·2H_2_O (0.76 g/l), ZnSO_4_ ·7 H_2_O (240 mg/l), Cu(NO_3_)_2_ ·3 H_2_O (40 mg/l), CoCl_2_·6 H_2_O (200 mg/l)]. For pH 6.0, the MOPS buffer was replaced with 100 mM MES (2-(*N*-morpholino)ethanesulfonic acid) and for pH 8.0, the buffer was 100 mM TAPS ([tris(hydroxymethyl)methylamino]propanesulfonic acid). All buffered media were adjusted for pH with KOH. Growth of the enriched culture of *Rhodoferax antarcticus* JLS was compared with culture of the type strain *Rhodoferax antarcticus* ANT.BR obtained from the American Type Culture Collection (ATCC 700587).

### DNA isolation and sequencing

Mat DNA was extracted by the PowerBiofilm kit (MO BIO). From each sample, approximately 20 mg material was extracted, and 0.75-1.50 µg DNA (measured by Qubit) was sent for sequencing. From lake water filters, 300-400 ng DNA was obtained by extraction using an MP FastDNA SPIN DNA kit (MP Biomedicals, CA) (Bielewicz *et al*. 2011). All metagenomic DNA was sequenced at DOE JGI (Joint Genome Institute Community Science Program award 1936). Shotgun metagenomic library construction and sequencing were carried out at JGI using standard protocols on the Illumina HiSeq 2500 platform. The raw reads were quality filtered using JGI standard protocols, and 1.2 Tb total sequences were obtained. Illumina reads were further processed using Trimmomatic (Bolger *et al*. 2014) to remove adapters and low-quality sequences. Trimmomatic counts the number of reads per FASTQ file, and it was used to determine the total reads per sample metagenome.

Cultured *Rhodoferax* biofilm DNA was isolated by the PowerBiofilm kit (MO BIO). Sequencing was performed by the Michigan State University Genomics Core. Libraries were prepared using the Illumina TruSeq Nano DNA Library Preparation Kit. Sequencing was performed on an Illumina MiSeq done in a 2×250bp format using an Illumina 500 cycle v2 reagent cartridge. The raw reads were quality filtered and 1 Tb total sequences were obtained.

### Identification of ARGs by ShortBRED

ShortBRED (Kaminski *et al*. 2015) is a pipeline that identifies protein family sequences by generating specific peptide markers from a database of interest, then screening the markers for specificity against the UniProt universal database of protein sequences (Bateman *et al*. 2021). The shortbred_identify command scans a reference database for protein family-specific peptide sequences. It was used to scan UniRef (the UniProt protein reference database) for sequences of the ARGs in version 3.0.7 of the Comprehensive Antibiotic Resistance Database (CARD) (Alcock *et al*. 2020). The minimum marker length accepted was 8 amino-acid residues, and the maximum length for combined markers for a given protein was 200. Markers that matched nonspecific genes in UniProt were discarded from the marker dataset.

The ShortBRED marker set was then used by the shortbred_quantify command to assign ARGs in the 150-kb reads from lake water and mat samples. From each target read, six reading frames were translated using tblastn. A marker hit required 95% match to the target. Sample class differences for ARG abundance were tested for significance (P ≤ 0.05) using the Kruskal-Wallis test.

### Taxa classification of metagenomic reads

The 150-kb sequence reads from mat and water metagenomes were assigned to taxonomic classes using the Kraken 2 classifier pipeline (Wood & Salzberg 2014, Wood *et al*. 2019). Kraken associates genomic *k*-mers (short sequence strings) in the reads with lowest common ancestor taxa. The read classifications were then used to calculate taxa percentages via the Bracken abundance estimator (Lu *et al*. 2017). The *k-*mer length was set at 150. For the Kraken 2-Bracken pipeline, the reference database used was the 5/17/2021 Standard Collection accessed at: https://benlangmead.github.io/aws-indexes/k2.

Eukaryotic taxa were further characterized using the pipeline EukDetect (Lind & Pollard 2021). This pipeline assigns reads to a database of 521,824 marker genes from 241 gene families, out of 3713 genomes and transcriptomes of fungi, protists and invertebrates.

For an alternative classifier, mat and water metagenomes were mapped to core taxonomic markers using the MetaPhlAn2 pipeline (Segata *et al*. 2012, Truong *et al*. 2015). MetaPhlAn2 assigns short reads to taxa using a set of marker genes identified from approximately 17,000 microbial reference genomes, primarily bacteria and archaea.

### Genome assembly by *breseq*

The pipeline *breseq* 0.35.6 (Deatherage & Barrick 2014) was used to assemble reads from two samples of the enrichment culture obtained from Mat-04. The reads were mapped to the *R. antarcticus* ANT.BRT (DSMZ24876) reference genome (Baker *et al*. 2017).

## RESULTS

### Taxa abundance of mat and water samples

Metagenomes were sequenced from three microbial sources: the water columns of Lake Fryxell (FRY-01, 02, 03) and Lake Bonney (BON-01, 02, 03), and the lift-off mat from the ice surface of Lake Fryxell (Mat 01 – 06). A total of 4 billion reads were sequenced, with a range of between 210 – 420 million reads per metagenome. The average read length was 147 bp, SD = 26.

From the three groups of samples, we characterized the taxa abundance using Kraken2/Bracken pipelines (Wood *et al*. 2019, Lu *et al*. 2017). The ice-surface mat samples (Mat-01 through Mat-06) showed substantial amounts of cyanobacteria, ranging from 9-60% of the total (**Fig. 2A**). The predominant orders of cyanobacteria were Oscillatoriales and Nostocales (**Fig. 2B**). High abundance of Oscillatoriales and Nostocales is consistent with previous reports for Fryxell mats obtained from benthic samples (Dillon *et al*. 2020). For Mat-01, 02, 03, 06 the cyanobacterial assignments were primarily Oscillatoriales and Nostocales. Mat-04 and Mat-05 however showed depletion of Oscillatoriales, with predominant abundance of Betaproteobacteria. Throughout the six mat samples, other taxa with significant abundance included Actinobacteria, Bacteroidetes, and Alphaproteobacteria, with smaller abundance of Planctomycetes, Bacteroidetes and Firmicutes. Thus the ice-trapped, air-exposed community showed a composition remarkably similar to that of the benthic mat samples from which lift-off mats originate (Dillon *et al*. 2020).

**Figure 2.**
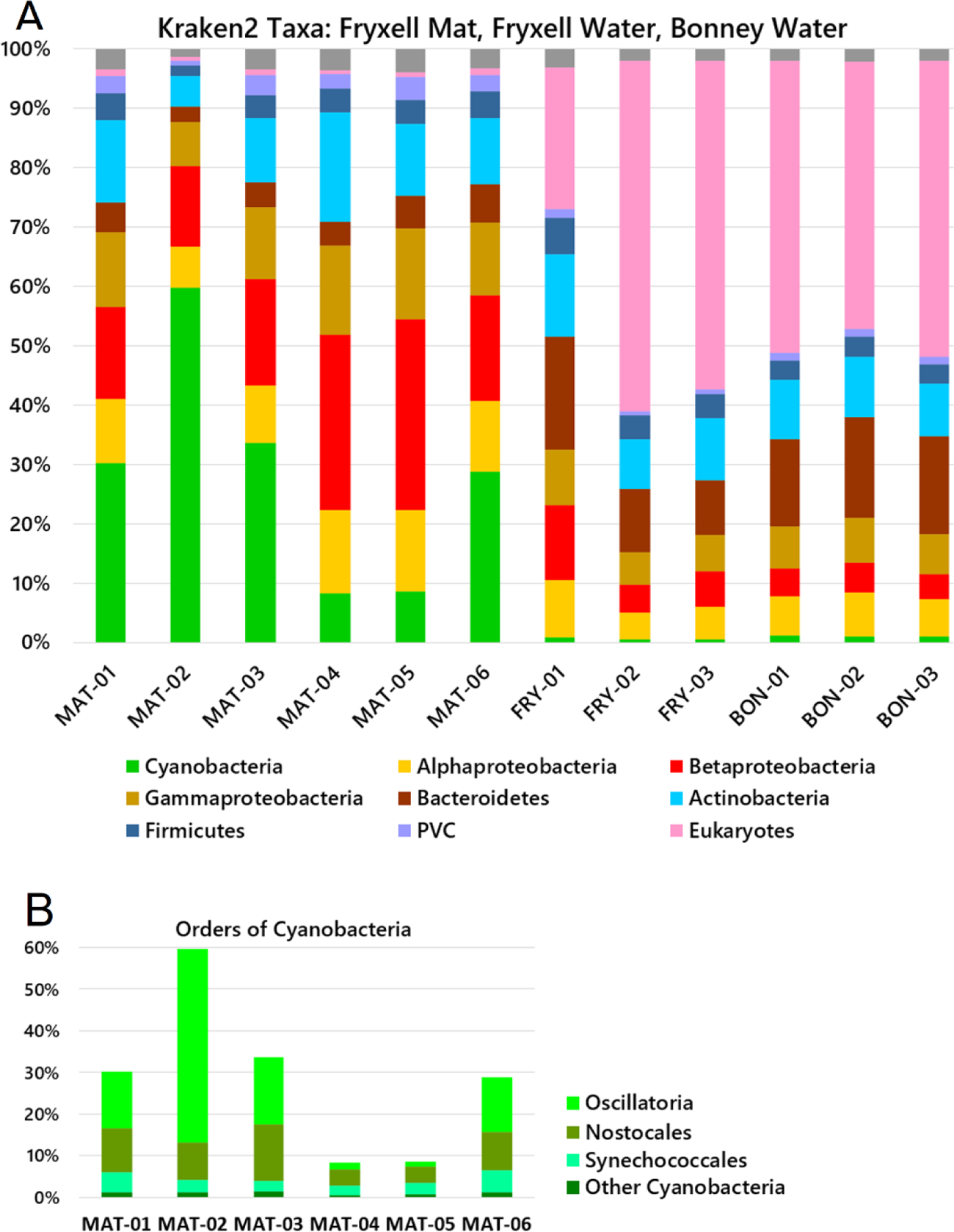
Microbial community composition of mat and water samples. Relative abundance of phyla was determined by read alignment to marker genes using Kraken2/Bracken pipeline. (**A**) Phyla and classes. (**B**) Orders of cyanobacteria in mat samples. MAT (Lake Fryxell, mat samples); FRY (Lake Fryxell, water samples); BON (Lake Bonney, water samples). MAT samples were from ice surface, unfiltered. The water samples were collected on a 0.45-µm filter. PVC=Planctomycetes-Verrucomicrobia-Chlamydiae

The filtered water from both lakes Fryxell and Bonney contained abundant bacterial taxa of Actinobacteria, Bacteroides, Betaproteobacteria, and Alphaproteobacteria, similar to previous reports of Fryxell water (Kwon *et al*. 2017) and other glacier-associated water bodies (Boetius *et al*. 2015). The water column showed almost no cyanobacteria but a high proportion of eukaryotes, in some cases more than 50% of the reads classified by Kraken2/Bracken. By contrast, the mat samples showed virtually no detectable eukaryotic DNA. The one-sided Mann-Whitney U test confirms that water from each lake contains more eukaryotic DNA than the mat samples (p = 0.01) and that the mat samples contain more cyanobacteria than the water microbiomes of either lake (p = 0.01).

To classify the eukaryotes, we used EukDetect (Lind & Pollard 2021) a pipeline that matches short read data to marker genes from fungal, protist and invertebrate genomes (**Fig. 3**; full output provided in **Table S1**). EukDetect identified the protist species *Geminigera cryophila* (van den Hoff *et al*. 2020), *Mesodinium rubrum* (Yih *et al*. 2004), *Nannochloropsis limnetica* (Kong *et al*. 2012), and *Chlamydomonas* sp. ICE-L (Lizotte *et al*. 1996). The two lakes showed similar taxa profiles, except that *Chlamydomonas* was found only in Lake Bonney. Mat samples had too few eukaryotic read counts to classify.

**Figure 3.**
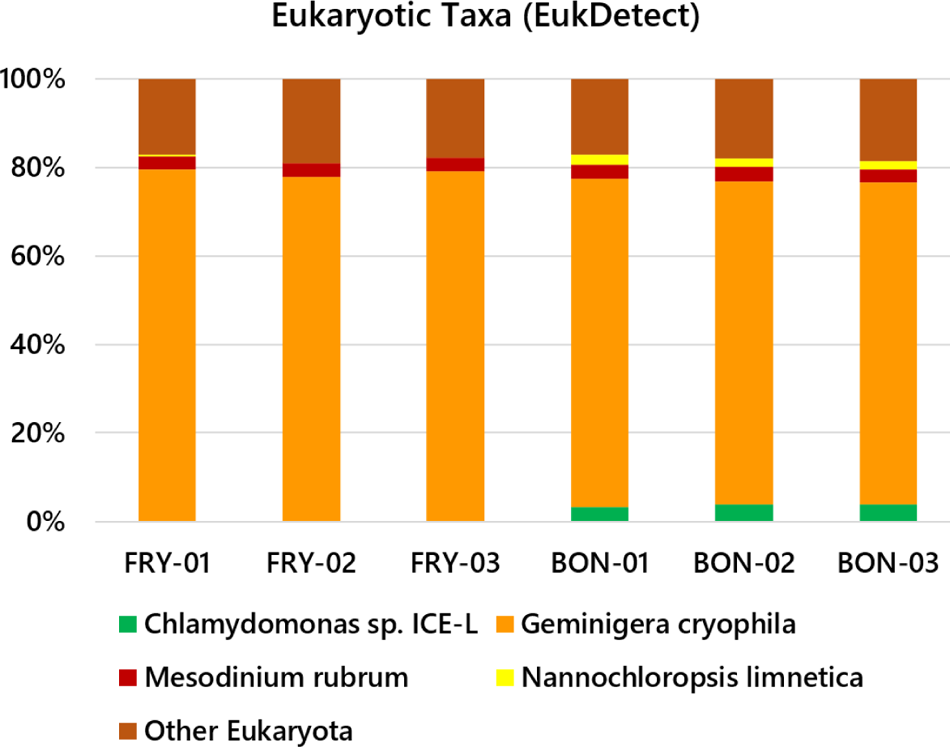
Eukaryotic taxa classified by EukDetect. Relative abundance of protist taxa were determined by read alignment to marker genes using EukDetect (Lind & Pollard 2021).

The validity of taxa classifier pipelines is highly dependent on their algorithm and taxa database. For comparison with Kraken2/Braken output, our mat and water reads were mapped to core taxonomic markers using an alternative pipeline, MetaPhlAn2 (Segata *et al*. 2012, Truong *et al*. 2015) (**Fig. 4**). Four of the six mat samples showed mainly cyanobacteria, consistent with longstanding reports of cyanobacterial mats (Taton *et al*. 2003, Hawes *et al*. 2013, Jungblut *et al*. 2016). Consistent with the Kraken analysis, Mat-04 and Mat-05 samples showed major amounts of Betaproteobacteria and Alphaproteobacteria, with depletion of cyanobacteria. For the filtered water samples, MetaPhlAn2 markers include limited content of Eukarya, and the pipeline did not find the eukaryotic taxa predicted by Kraken2 (**Fig. 2**). The filtered water from both lakes showed mainly Actinobacteria, with some Betaproteobacteria. Overall MetaPhlAn2 appeared less effective than Kraken 2/Bracken at classifying our samples.

**Figure 4.**
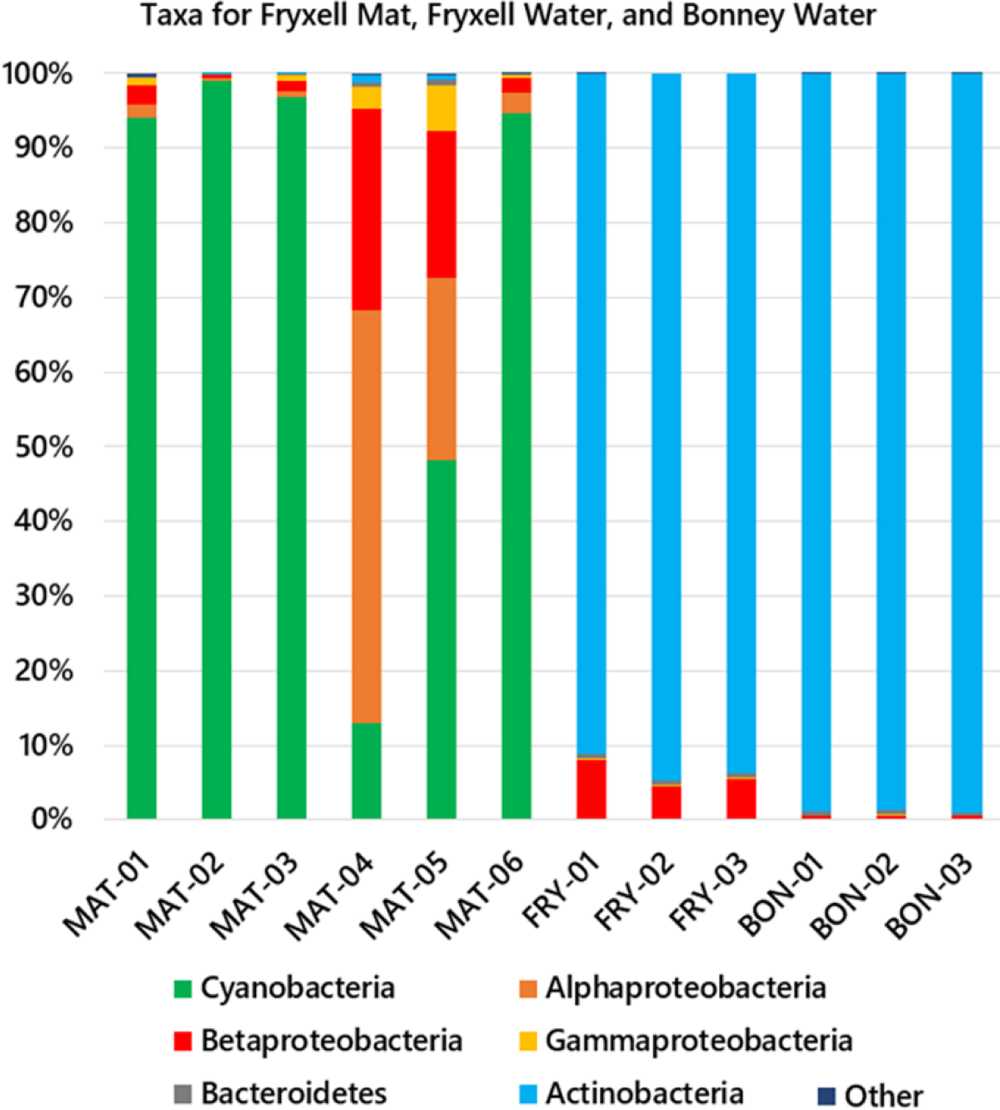
Microbial community composition via MetaPhlAn2. Relative abundance of class-level taxa was determined by read alignment to marker genes using MetaPhlAn2 (Segata *et al*. 2012, Truong *et al*. 2015). Samples were as indicated for Figure 2.

### ARG composition characterized using ShortBRED

Gene sequences encoding various forms of antibiotic resistance, including genomic loci as well as mobile elements, are collected in the Comprehensive Antibiotic Resistance Database (CARD) (Alcock *et al*. 2020). We used the ShortBRED pipeline to identify ARGs in our samples, based on a set of marker peptides matching CARD sequences. The percentage of reads from each sample that matched ARGs ranged from 0.0001-0.0006% (**Fig. 5**). The overall ARG abundances were similar amongst the three sample classes (Fryxell mat, Fryxell water, Bonney water).

**Figure 5.**
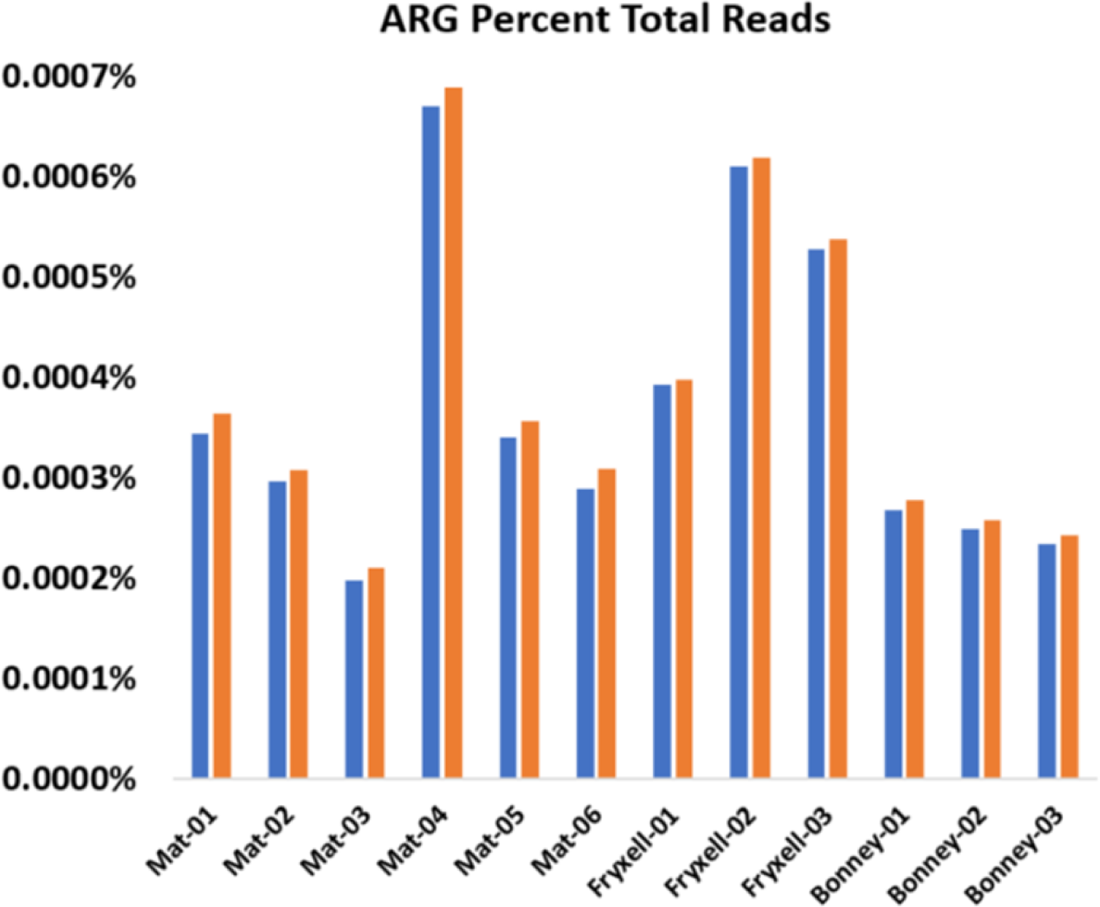
ARG abundance in lake samples. Percentage of reads matched to ShortBRED markers. Total reads were counted using Trimmomatic. Bars indicate total ARG hits per sample (blue) and top 60 ranked ARG hits (orange).

The specific ARGs identified by ShortBRED were sorted by abundance across samples (**Table 2**; Supplementary file **Table S2**). Eight of the top 20 sorted ARGs showed a difference among the three sample classes (Kruskal-Wallis test, P ≤ 0.05); of these results, 7 of 8 are significant. Of the top-ranked ARGs, BUT-1 and vancomycin-resistance genes *vanYA*, *vanTG* and *vanI* appeared mainly in the water column. BUT-1 is class-C beta-lactamase reported in *Buttiauxella agrestis*, an environmental Gammaproteobacterium found in environmental water sources (Fihman *et al*. 2002, Nakamichi *et al*. 2021). The vancomycin resistance genes occur in *Enterococcus* and other Gram-positives (Courvalin 2006). By contrast, the beta-lactamase AAC(3)-Ia (Riccio *et al*. 2003) occurred only in the mat samples. AAC(3)-Ia is encoded on *Pseudomonas aeruginosa* integrons as well as in other Proteobacteria. Other ARGs showed possible differences in prevalence between Lake Bonney and Lake Fryxell water; for example, the Streptomyces-associated beta-lactamase AAC(3)-VIIa (López-Cabrera *et al*. 1989) was more abundant in Bonney water whereas *vanYA* was more abundant in Fryxell water.

**Table 2.**
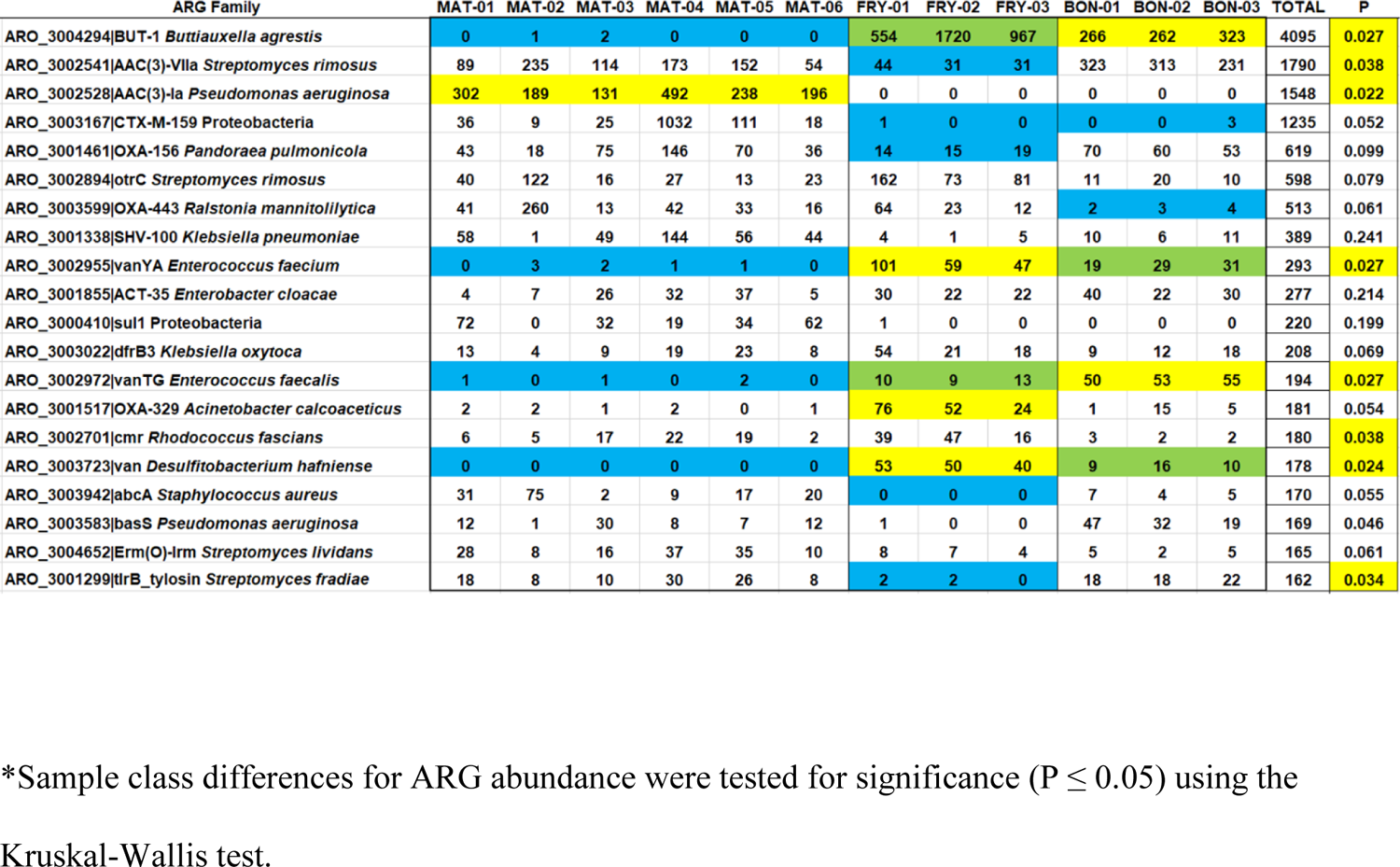
Mat and water antibiotic resistance genes (ARGs) identified by ShortBRED marker peptides*

Overall, the three sample classes each had distinctive ARG-abundance signatures, as indicated by Kruskal-Wallis test. Ten of the top 20 ranked ARGs were associated with Proteobacteria, which roughly corresponds with the proportion of bacterial taxa abundances predicted by Kraken2/Bracken (**Fig. 2**).

### Enrichment of Mat-forming *Rhodoferax antarcticus*

The relatively high proportions of Betaproteobacteria and Alphaproteobacteria in MAT-04 differed from that of other samples, and the color of the sample material was more red than green. We cultured the various samples anaerobically (in closed tubes) at 10°C with illumination. From MAT-04 red spots of biofilm were obtained. There was no evidence of planktonic growth (cloudiness) or cell motility within the culture medium, but the organism grew as a mat, forming red blobs with a “red nose” appearance. In addition the culture extended as a red film-like mat growing along the inside of the culture tube (**Fig. 6A**). It was not possible to obtain isolated colonies, but serial culture of the “red noses” showed consistent biofilm formation and Gram-stain morphology (**Fig. 6B**). This finding is consistent with an earlier report that the Fryxell sedimentary biofilm includes a layer of phototrophic purple bacteria beneath the cyanobacterial layer (Buffan-Dubau *et al*. 2001). Ours is the first report of such biofilm cultured from material at the ice surface, rather than from its inferred origin in the sediment at the bottom of the lake.

**Figure 6.**
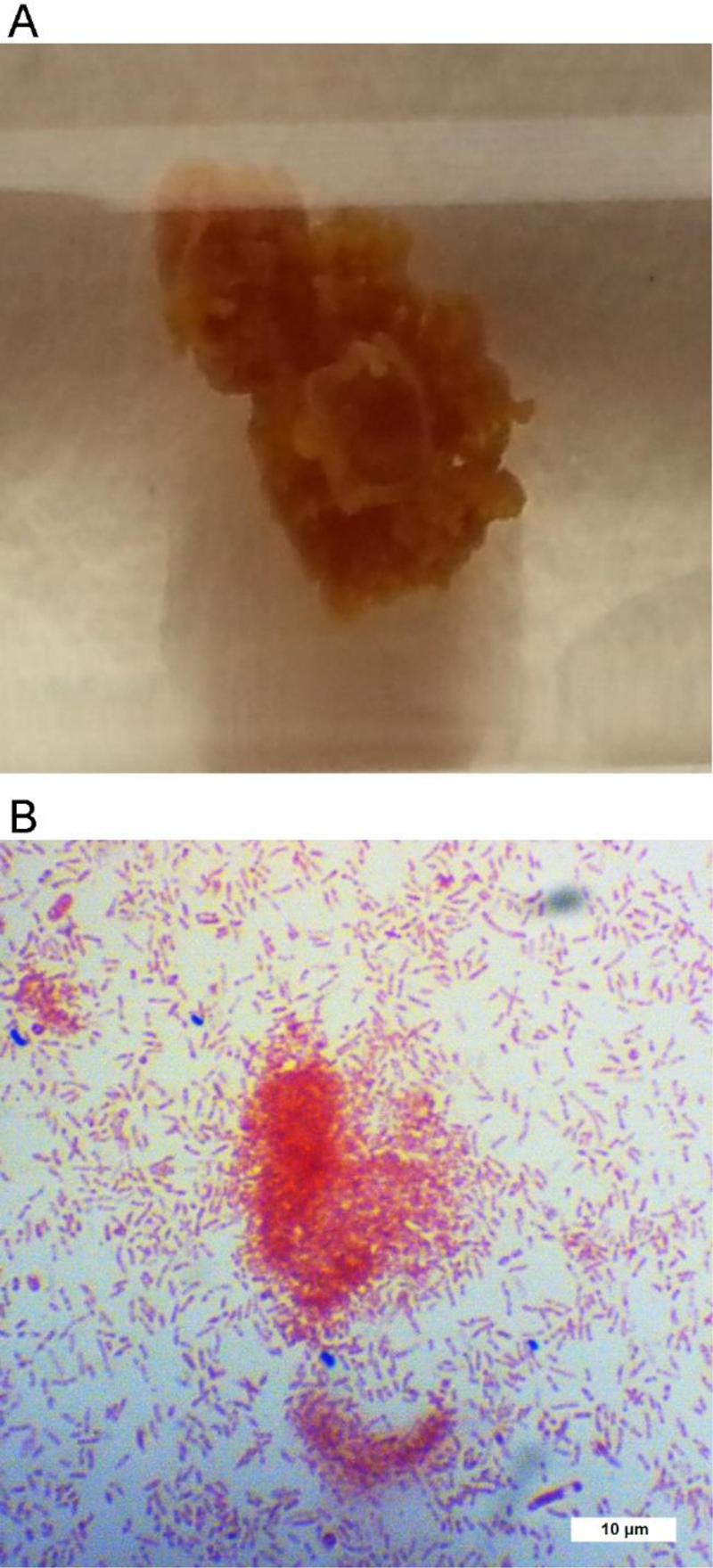
Mat-forming enrichment of Mat-04 *R. antarcticus* JLS. **(A)** Biofilm of cultured *R. antarcticus* within closed tube of medium, showing globular form of growth. Initial Mat-04 sample was subcultured serially three times in PM medium, in horizontal closed tubes at 10°C with 30 umoles/m^2^s illumination. **(B)** Gram stain of culture, 1000X with oil immersion. Courtesy of Emma Stuart-Bates.

The MAT-04 biofilm culture was subjected to genome analysis. DNA was sequenced and short reads were used to BLAST the NCBI database. The predominant hits were to the genome of *Rhodoferax antarcticus* ANT.BRT, a Betaproteobacterium originally isolated from a saltwater pond at Cape Royds, Ross Island, Antarctica (Madigan *et al*. 2000, Baker *et al*. 2017). Kraken 2/Bracken analysis of sequenced reads agreed with this classification, assigning the samples as 95% *R. antarcticus* with about 1% *Pseudomonas*.

We used the *breseq* pipeline (Deatherage & Barrick 2014) to assemble reads from two non-axenic samples (7-RN1 and 8-RN2) whose DNA sequence reads were assembled to the *R. antarcticus* ANT.BRT reference genome (Supplementary file, **Table S3** and **S4**). These tables present “mutations,” that is, all differences from the ANT.BR reference sequence that show 100% read coverage. The two samples, 7-RN1 and 8-RN2, showed 276 and 271 sequence differences respectively, in their main chromosome compared with that of the reference genome, length 3,809,266 bp (Baker *et al*. 2017). The sample genomes showed > 99.99% identity with the main chromosome and > 99% identity with one plasmid in the reference. A large part of the observed sequence differences from reference were silent mutations. These observations confirm the close relatedness of our culture to the published strain.

The *R. antarcticus* enrichment culture (designated *R. antarcticus* JLS) was tested for ARG abundance using ShortBRED (**Table 3**). More than thirty ARGs showed hits, but nearly all were associated with *Pseudomonas aeruginosa*. By contrast, no ARGs matched those from CARD database that were associated with *Rhodoferax*.

**Table 3.**
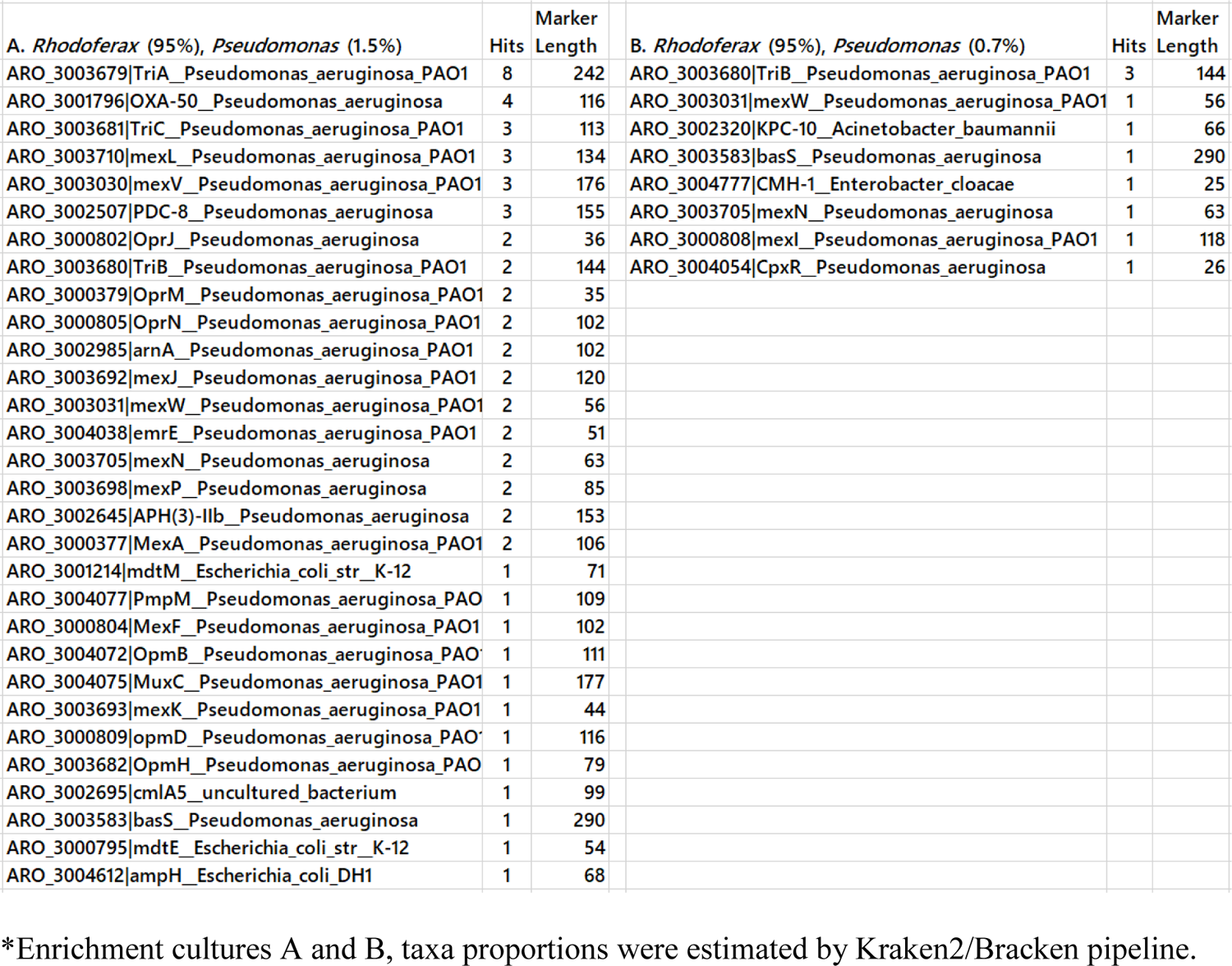
*Rhodoferax* enrichment culture ARGs identified by ShortBRED*

### Characterization of the mat-forming *R. antarcticus* JLS

The *R. antarcticus* JLS biofilm was further tested for growth range of pH and salinity. The biofilm was subcultured anaerobically with illumination, in a malate-succinate *Rhodoferax* medium modified from that of references (Madigan *et al*. 2000, Tayeh & Madigan 1987) as described under Methods (**Fig. 7A, B, C**). After 45 days, at pH 7, the bacteria formed globules as well as a red coating along the glass. At pH 6, spots of biofilm grew slowly, and at pH 8 little growth was seen. Growth was also tested for cultures buffered at pH 7 with NaCl amendment (**Fig. 7D, E, F**). The fullest growth was seen in the absence of NaCl amendment (the core medium contains approximately 10 mM Na^+^ ions). Less growth was seen with added NaCl (23 mM or 46 mM).

**Figure 7.**
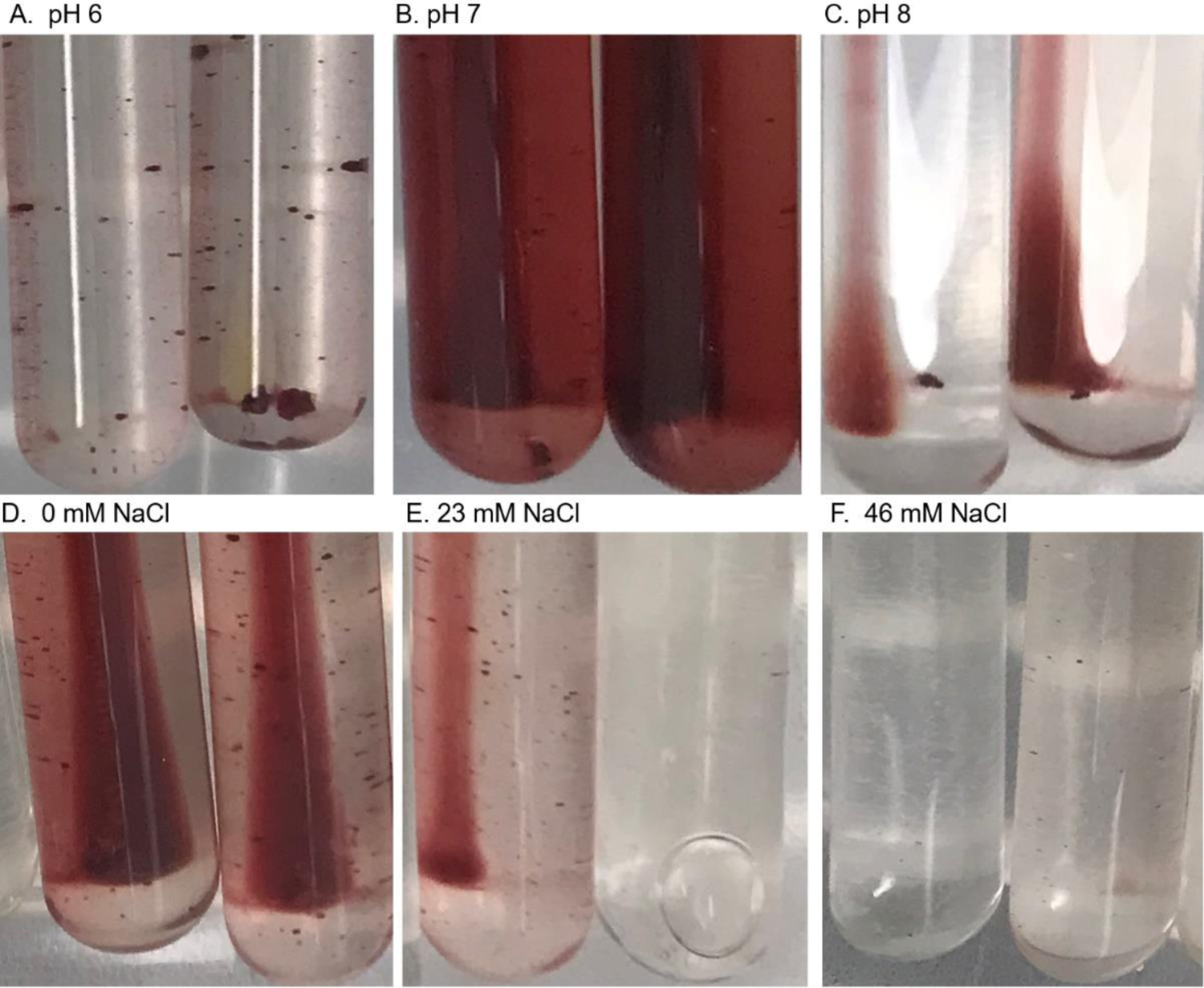
pH and NaCl dependence of *R. antarcticus* JLS. **(A, B, C)** Medium PM contained **(A)** 100 mM MES pH 6; **(B)** 100 mM MOPS pH 7; **(C)** 100 mM TAPS pH 8. Cultures were incubated 45 days. (**D, E, F)** Medium PM was amended with **(D)** no NaCl added; **(E)** 23 mM NaCl; **(F)** 46 mM NaCl. Cultures were incubated 45 days.

Despite the high sequence identity, the *R. antarcticus* JLS enrichment culture differed from the *R. antarcticus* ANT.BR strain (Madigan *et al*. 2000) in its growth phenotype, under all conditions of pH and NaCl concentration that were tested for both strains. Unlike the motile, planktonic *R. antarcticus* ANT.BR, our culture showed no motility and little sign of planktonic single-celled growth. Instead, the subcultured material grew entirely as a biofilm, in globular spots and as a film-like growth along the interior surface of the glass tube.

The type strain ANT.BR was cultured at the same time, under all conditions. The original strain never formed a biofilm; it appeared planktonic and motile under all conditions of pH and NaCl concentration.

## DISCUSSION

Identification of antibiotic resistance genes from relatively pristine environments is of interest for several reasons. Long before human introduction of high-dose antibiotics, environmental bacteria evolved multidrug pumps to efflux toxic products of their own metabolism, as well as antimicrobial substances produced by their competitors (Allen *et al*. 2010, Wright 2019). Many antibiotics possess signalling capabilities and other unknown functions. Phylogeny dates the origin of beta-lactamases to hundreds of millions of years ago. Vancomycin resistance genes are found in 30,000-year-old permafrost (Dcosta *et al*. 2011). But the specific kinds of ARGs found in environmental sources may differ from those prevalent in human microbiomes, those conferring resistance to the drugs we depend on for therapy (Zeng *et al*. 2019).

In our Taylor Valley lake metagenomes, the top-scoring ARG was BUT-1, a cephalosporinase previously found in a clinical isolate of *Buttauxiella* (Fihman *et al*. 2002). Two other ARGs of clinical origin (*vanYA*, *vanTG*) encode vancomycin resistance components in *Enterococcus* (Courvalin 2006, Boyd *et al*. 2006). The finding of clinical ARGs in Antarctic water bodies is concerning. The rest of the top 20 ARGs we found appear common in environmental organisms. Eight were beta-lactamases, which are commonly found in environmental organisms but can also be readily transferred between environmental and pathogenic strains (Hooban *et al*. 2020).

In polar regions, previous metagenomic studies reveal a range of naturally occurring ARGs. Surface soils of the Antarctic Mackay Glacier region show ARGs encoding multidrug pumps, beta-lactamases and aminoglycoside inactivators, largely associated with Gram-negative bacteria (van Goethem *et al*. 2018). A study of Tibetan soils showed a number of abundant ARGs, most notably encoding vancomycin resistance (Bo Li *et al*. 2020). Our study of ARGs from Antarctic lakes adds to this picture, showing that both water and microbial mat sources contain familiar ARGs, and likely contain others not yet discovered in Antarctic strains.

It was interesting to compare the total ARG abundance of our Antarctic lake samples with those of temperate-zone Ohio rural water bodies with moderate human inputs, in a study where samples were prepared by the same method and analyzed using the same ShortBRED marker set (Murphy *et al*. 2021). The overall ARG abundance is about the same for the two environmental sources, except for river samples obtained just downstream of a wastewater effluent pipe, where the total ARG prevalence is increased approximately five-fold. This result is consistent with a model that natural microbial communities generally harbor a small balanced prevalence of ARGs, which gets amplified by concentrated human input. Note however that a database such as CARD can only represent a fraction of the actual ARGs out there; new drug resistance families and related mobility agents are continually discovered.

The mat lift-off samples we obtained from the ice surface showed a range of DNA taxa consistent with those reported for samples obtained from benthic mats (Dillon *et al*. 2020). Given that our collected mat organisms had survived several years within ice, followed by air dessication and prolonged UV radiation exposure, it is impressive how much of the Antarctic mat microbiomes retain viability with intact DNA. Even ciliated protists and invertebrate worms were obtained alive from samples cultured after months of storage at −80°C (not shown). It is likely that some of these organisms are psychrophiles that continue to grow within the ice (Boetius *et al*. 2015).

Previous studies emphasize the cyanobacterial content of lift-off mats, primarily Oscillatoriales genera such as *Microcoleus*, as well as *Nostoc* (Taton *et al*. 2003, Jungblut *et al*. 2016). While most of our lift-off samples showed abundance of cyanobacteria, one sample yielded cultures from which the majority of reads matched *R. antarcticus*. The finding of mat samples enriched for *R. antarcticus* indicates that portions of the lower layer of *Rhodoferax* mat (Buffan-Dubau *et al*. 2001), along with the cyanobacterial upper layer, can break off and form lift-off patches that emerge from the ice. Our culture of *R. antarcticus* (*R. antarcticus* JLS) obtained from Lake Fryxell showed mat-forming morphology very different from the motile single cells of *R. antarcticus* ANT.BR isolated from Cape Royds (Madigan *et al*. 2000, Baker *et al*. 2017). Despite the high genetic similarity, our cultured organism appears to represent a novel ecotype of *R. antarcticus*.

Our genomic reads from *R. antarcticus* JLS showed no matches to our ShortBRED antibiotic resistance markers, although the reference genome does indeed include various resistance genes including numerous RND and MFS transporters as well as multidrug efflux components. Thus, many naturally-occurring ARGs are likely to be missed by standard marker searches.

The prevalence of eukaryotic sequences in the lake water metagenomes is consistent with previous reports that protists play important roles in the Taylor Valley lake communities (Li *et al*. 2016, Bielewicz *et al*. 2011, Glatz *et al*. 2006, Lizotte *et al*. 1996). Microbial eukaryotes including phototrophs and mixotrophs provide prominent functions in the lake ecology (Li *et al*. 2016). In our data, the eukaryotic community of Lakes Fryxell and Bonney showed three major taxa in common: *Gemingera cryophila*, *Mesodinium rubrum*, and *Nannochloropsis limnetica*. *G. cryophila* is a mixotrophic cryptophyte that feeds on bacteria but also conducts photosynthesis as a secondary endosymbiont alga (van den Hoff *et al*. 2020). *M. rubrum* is a ciliate that consumes cryptophytes but also uses the prey chloroplasts to conduct photosynthesis (kleptoplasty) (Yih *et al*. 2004). *N. limnetica* is a heterokont alga, with red alga-derived chloroplasts, primarily a phototroph (Kong *et al*. 2012). Our Lake Bonney samples also showed sequences from *Chlamydomonas*, a green alga that dominates some parts of the Lake Bonney water column (Lizotte *et al*. 1996).

We note that in the water samples, smaller aquatic phototrophs were likely missed by the 0.45-µm filter; 0.2-µm filters would have been preferable but were not available in the field at the time. Even 0.2-µm filters miss important microbial community members (Brown *et al*. 2015). The mat samples however had no filtration, so a broader spectrum of cell sizes was captured.

It is interesting that the Lake Fryxell water shows mainly eukaryotic phototrophs whereas the mat shows mainly cyanobacteria and proteobacterial phototrophs. The mat bacteria are likely to survive a wider range of light and temperature conditions than the eukaryotes. From the standpoint of drug resistance, cyanobacteria are more likely than eukaryotes to harbor and transfer ARGs of potential bacterial pathogens. Nevertheless, protists can regulate bacterial ARG composition in terrestrial communities (Nguyen *et al*. 2020), so this factor may be of interest when assessing the lake water ARG pools.

## Acknowledgments and Funding

Fieldwork was supported by the National Science Foundation award OPP-1637708 to RM. For metagenome sequencing we thank the U.S. Department of Energy Joint Genome Institute, a DOE Office of Science User Facility, supported by the Office of Science of the U.S. Department of Energy Contract No. DE-AC02-05CH11231, JGI Metagenome award 1936 to JLS. Bioinformatic analysis and biofilm culture and sequencing were supported by the National Science Foundation award MCB-1923077 to JLS. We thank Caroline Harwood for guidance regarding culture of the mat biofilm, and Emma Stuart-Bates for the biofilm culture micrograph. We especially thank Darcy Blankenhorn for expert technical support.

## Author Contributions Statement

ST designed the bioinformatic analysis of ARGs and drafted the manuscript. GH characterized the *Rhodoferax* enrichment culture and contributed to the manuscript. DB ran the bioinformatic pipelines and performed statistics. RM led the field expedition and collected water samples. JLS conceived the central concept, collected the mat samples and isolated DNA, mentored students, and completed the manuscript.

## Availability of data and materials

Sequence read files from lake water and mat samples are deposited at NCBI under SRA accession numbers SRP104818, SRP104821, SRP098041, SRP098040, SRP098042, SRP104817, SRP098044, SRP104822, SRP098050, SRP104819, SRP104820, SRP104823. Sequence read files from cultured *Rhodoferax antarcticus* JLS are deposited at NCBI under SRA accession number PRJNA736311.

